# Evaluating Mendelian nephrotic syndrome genes for evidence of risk alleles or oligogenicity that explain heritability

**DOI:** 10.1101/071639

**Authors:** Brendan Crawford, Christopher E. Gillies, Catherine C. Robertson, Matthias Kretzler, Edgar Otto, Virginia Vega-Warner, Hyun Min Kang, Matthew G. Sampson

**Affiliations:** Department of Pediatrics, University of Michigan Medical School; Department of Internal Medicine, University of Michigan Medical School; Department of Biostatistics, University of Michigan School of Public Health

**Keywords:** Nephrotic Syndrome, Variant, Mendelian;, Oligogenicity, Risk allele

## Abstract

**Background:** More than 30 genes can harbor rare exonic variants sufficient to cause nephrotic syndrome (NS), and the number of genes implicated in monogenic NS continues to grow. However, outside the first year of life, the majority of affected patients, particularly in ancestrally mixed populations, do not have a known monogenic form of NS. Even in those children classified with a monogenic form of NS, there is phenotypic heterogeneity. Thus, we have only discovered a fraction of the heritability of NS – the underlying genetic factors contributing to phenotypic variation. Part of the “missing heritability” for NS has been posited to be explained by patients harboring coding variants across one or more previously implicated NS genes, insufficient to cause NS in a classical Mendelian manner, but that nonetheless impact protein function enough to cause disease. However, systematic evaluation in patients with NS for rare or low-frequency risk alleles within single genes, or in combination across genes (“oligogenicity”), has not been reported.

**Objective:** To determine whether, as compared to a reference population, patients with NS have either a significantly increased burden of protein-altering variants (“risk-alleles”), or unique combination of them (“oligogenicity”), in a set of 21 genes implicated in Mendelian forms of NS.

**Methods:** In 303 patients with NS enrolled in the Nephrotic Syndrome Study Network (NEPTUNE), we performed targeted amplification paired with next-generation sequencing of 21 genes implicated in monogenic NS. We created a high-quality variant call set and compared it to a variant call set of the same genes in a reference population composed of 2535 individuals from Phase 3 of 1000 Genomes Project. We created both a “stringent” and “relaxed” pathogenicity filtering pipeline, applied them to both cohorts, and computed the (1) burden of variants in the entire gene set per cohort, (2) burden of variants in the entire gene set per individual, (3) burden of variants within a single gene per cohort, and (4) unique combinations of variants across two or more genes per cohort.

**Results:** With few exceptions when using the relaxed filter, and which are likely the result of confounding by population stratification, NS patients did not have significantly increased burden of variants in Mendelian NS genes in comparison to a reference cohort, nor was there any evidence of oligogenicity. This was true when using both the relaxed and stringent variant pathogenicity filter.

**Conclusion:** In our study, the burden or particular combinations of low-frequency or rare protein altering variants in previously implicated Mendelian NS genes cohort does not significantly differ between North American patients with NS and a reference population. Studies in larger independent cohorts or meta-analyses are needed to assess generalizability of our discoveries and also address whether there is in fact small but significant enrichment of risk alleles or oligogenicity in NS cases undetectable with this current sample size. It is still possible that rare protein altering variants in these genes, insufficient to cause Mendelian disease, still contribute to NS as risk alleles and/or via oligogenicity. However, we suggest that more accurate bioinformatic analyses and the incorporation of functional assays would be necessary to identify bona fide instances of this form of genetic architecture as a contributor to the heritability of NS.

## INTRODUCTION

Nephrotic syndrome (NS) is a rare disorder of glomerular filtration barrier damage, clinically characterized by proteinuria, hypoalbuminemia, and edema [1]. In 1998, pedigree analysis of familial NS identified disease-causing variants within *NPHS1*, identifying it as a single-gene (monogenic) cause of NS [2]. Since that time, genetic discovery in familial NS have identified over 30 genes implicated in monogenic forms of disease (reviewed in [3, 4]). The discovery of these genes has shed insight into the molecular mechanisms underlying NS (in particular implicating the podocyte as the major cell affected by NS). From a clinical perspective, genotype-phenotype studies suggest patients classified with monogenic NS are resistant to immunosuppressive therapy and have faster renal functional decline, but do not develop disease recurrence following transplant [5-9].

Yet despite the large number of genes already implicated, the majority of patients with NS (outside of those with congenital NS) have not been classified with a known Mendelian form of NS. The two largest published NS cohorts sequenced between 1-31 genes in 1174 patients [10] and 27 genes in 2016 patients [11]. The prevalence of monogenic NS in these two cohorts, enriched for SRNS and familial disease, was 23% and 30%, respectively.

While additional monogenic forms of NS continue to be discovered [12, 13], the percentage of NS patients in a population who will ultimately be classified as monogenic is unclear. This may particularly true for patients from more genetically diverse populations. For example, we recently performed targeted sequencing of 21 known NS genes in a North American population-based cohort of biopsied NS patients enrolled in the Nephrotic Syndrome Study Network (NEPTUNE). Using stringent filtering criteria, we classified only 3% of patients with a monogenic cause of their NS [14]. When restricting this analysis solely to children with FSGS or MCD, this prevalence was 5.5%.

Altogether, these prevalence estimates raise the question of whether alternative forms of genetic variation within these known NS genes may explain a portion of its heritability. Given their importance in the functioning of the glomerular filtration barrier, it has been an attractive hypothesis that protein altering variants within known monogenic NS genes, insufficient to cause monogenic disease themselves, could lead to disease. One alternative form proposed suggests that NS can be caused by an individual harboring protein-altering variants across more than one known monogenic NS gene (“oligogenicity”). For example, there is one report of a child with SRNS who progressed to end-stage renal disease (ESRD) by 8 years of age who had heterozygous variants in *NPSH2* and *CD2AP*[15]. While the child’s mother had a single *NPHS2* variant and father had a single *CD2AP* variant, neither parent had proteinuria while both had normal renal function. In another example, a child with Pierson syndrome had heterozygous variants in *LAMB2* and *NPHS1,*with the authors suggesting that each contributed to the patient’s disease [16]. In yet another example, a child with a pathogenic *WT1* variant and early-onset disease also had a single *NPHS1* variant; however, it is unclear if the *NPHS1* variant had a modifying effect on the phenotype [17]. Five individuals have been reported with two *NPHS1*variants and an *NPHS2*variant, suggesting possible phenotypic modifier effects (“tri-allelism”) [18, 19]. More recently, in a recent panel sequencing study of 37 SRNS patients, 22% of the 14 individuals with monogenic NS has a putatively pathogenic variant in an additional gene [16]. Oligogenicity has also been suggested in the pathogenesis of a number of other kidney diseases, including Bardet-Biedl syndrome, nephronopthisis, and autosomal dominant polycystic kidney disease [20-22].

Another alternative form of heritability involving Mendelian NS genes could from result rare or low-frequency protein-altering variants in a single gene that are insufficient to cause disease alone, but nonetheless greatly increase risk of NS. We use the term “risk alleles” to describe this potential form of NS-associated genetic architecture within implicated genes. Perhaps the first example of this comes from the pR229Q variant in podocin (*NPHS2*), which is present in ~3.6% of the European population, but contributes to monogenic NS when found in compound heterozygosity with specific *NPHS2*protein altering variants in trans [23]. More recently, a study applied rare variant association testing to exon sequencing data of 3000 genes in 214 Europeans with FSGS. They discovered protein-altering variants (including those within *APOL1*and *COL4A4)*, present rarely in the population, which were associated with significantly increased risk of disease [13].

Despite these publications, systematic evaluation of whether oligogenicity across Mendelian NS genes, or risk alleles within them, explain a portion of heritability of NS has not been reported. To address this question, we analyzed sequence data from 21 implicated Mendelian NS genes in 303 patients with NS enrolled in the Nephrotic Syndrome Study Network (NEPTUNE) cohort (who were previously determined to not have monogenic NS due to variants in these genes) [14] and ~2500 subjects from Phase 3 of the 1000 Genome project (1000G) [24]. We then tested the hypothesis that, as compared to the reference population, patients with NS were enriched for oligogenicity and/or risk alleles within this geneset.

## CONCISE METHODS

### Study Participants

The Nephrotic Syndrome Study Network is a longitudinal observational cohort study, composed of 23 centers across the US and Canada, recruiting participants across the age-range at their first clinically indicated biopsy for suspicion of having minimal change disease, FSGS, or membranous nephropathy [25, 26]. Participants were recruited without selection for positive family history of NS, age of onset, or response to therapy. 312 participants with available DNA for sequencing were previously sequenced across these 21 genes to determine the prevalence of monogenic NS in this cohort [14]. We excluded the nine individuals already classified with monogenic NS, resulting in 303 participants included in this study (**Table 1**). Because rare, protein-altering variants also exist in the general population in people without NS, we also studied the prevalence of these variants in the 1000 Genomes Project as the reference population [24]. These patients do not have any clinical phenotype data, but were not selected based on any clinical criteria. Given that NS is a rare disease, we assume that its prevalence was rare in this reference cohort. However, we did exclude 37 individuals with variants qualifying as monogenic disease, yielding 2498 individuals in reference population available for comparison.

**Table 1.** Demographic and clinical characteristics of 303 patients with NS in NEPTUNE cohort queried in this study. * 7 patients did not have age of disease onset reported. ** 1 patient did not have histopathologic diagnosis reported.

|  | NEPTUNE cohort (n= 303) |
| --- | --- |
| Pediatric Onset* | 98 (32%) |
| Median Age of Disease Onset(years) [IQR]* | 32 [15-52] |
| Pediatric | 10 [4-14] |
| Adult | 43 [31-58] |
| Male Gender | 182 (60%) |
| Race by patient report |  |
| White | 165 (54%) |
| Black | 73 (24%) |
| Asian & Pacific Islander | 37 (12%) |
| Multiple races | 14 (5%) |
| Not Reported | 14 (5%) |
| Histopathologic Diagnosis** |  |
| MCD | 62 (20%) |
| FSGS | 100 (33%) |
| MN | 50 (17%) |
| Other | 90 (30%) |
| Remission History |  |
| Achieved complete remission | 140 (46%) |
| Achieved partial but not complete remission | 96 (32%) |
| Never achieved partial or complete remission | 67 (22%) |

### Targeted Sequencing of 21 Genes and Sequence Data Analysis

For cases, microfluidic PCR paired with next generation sequencing of all exons of 21 monogenic NS genes was performed (**Table 2**), followed by sequence alignment, variant calling and functional annotation of variants was performed as previously described [14]. Variant data for these same genes was previously generated in the 1000G cohort and was available for analysis [14, 27].

**Table 2.** Mendelian “monogenic” nephrotic syndrome genes targeted for sequencing

| **Dominant** | **Recessive** |
| --- | --- |
| *ACTN4* | *COQ2* |
| *CD2AP* | *COQ6* |
| *CFH* | *CUBN* |
| *INF2* | *ITGA3* |
| *LMX1B* | *LAMB2* |
| *TRPC6* | *MYO1E* |
| *WT1* | *NEIL1* |
|  | *NPHS1* |
|  | *NPHS2* |
|  | *PDSS2* |
|  | *PLCE1* |
|  | *PTPRO* |
|  | *SCARB2* |
|  | *SMARCAL1* |

### Defining risk alleles and oligogenicity

In this study we defined “risk allele” as a variant in a previously implicated NS gene that is found in less than 1% of the population, and which is insufficient alone to cause NS. We defined “oligogenicity” as a collection of risk alleles in >1 previously implicated NS gene in an individual not classified with known monogenic NS. We give examples of each in **Table 3**. Risk alleles could include (1) a heterozygous variant in a recessive NS gene that would be sufficient to cause Mendelian NS if found homozygously or in compound heterozygosity with a causal allele *in trans* for the same gene, or (2) a heterozygous variant in a dominant NS gene which is insufficient alone to cause NS alone based on a stringent Mendelian pathogenicity filter.

**Table 3.** Hypothetical forms for complex heritability from Mendelian NS genes

| **Form of Inheritance** | **Description** | **Example** |
| --- | --- | --- |
| “Risk Allele” | Heterozygosity for a variant in a recessive NS gene OR heterozygosity for a variant in a dominant NS gene that does not fulfill characteristics of a Mendelian allele, whether in terms of allele frequency, conservation, or predicted impact on function | Heterozygosity for *MYO1E* p.I384T |
| “Oligogenicity” | Heterozygosity for a single pathogenic allele in multiple genes. Each variant allele is alone solely insufficient to cause phenotype but combined effect contributes to phenotype | Heterozygous for *CD2AP* p.M496I, AND heterozygous for *NPHS2* p.A208T |

To identify variants in NS cases and the reference population that qualified as a risk allele alone, or contributing to an oligogenic case, we constructed both a “stringent” and “relaxed” filter. **Table 4** displays the parameters for the stringent and relaxed pathogenicity filter for loss of function and missense variants. The Exome Aggregation Consortium (ExAC) contains sequences of around 60,000 individuals, and is currently an ideal resource for determining allele frequencies in populations [28]. However, ExAc contains the sequences of individuals for 1000 Genomes Project, the reference population for this study. Thus, we did not use the ExAC database; instead, the Exome Variant Server (EVS), paired with 1000G local ancestry-based frequencies, was used to incorporate allele frequencies in the filter, as previously published [14].

**Table 4.**
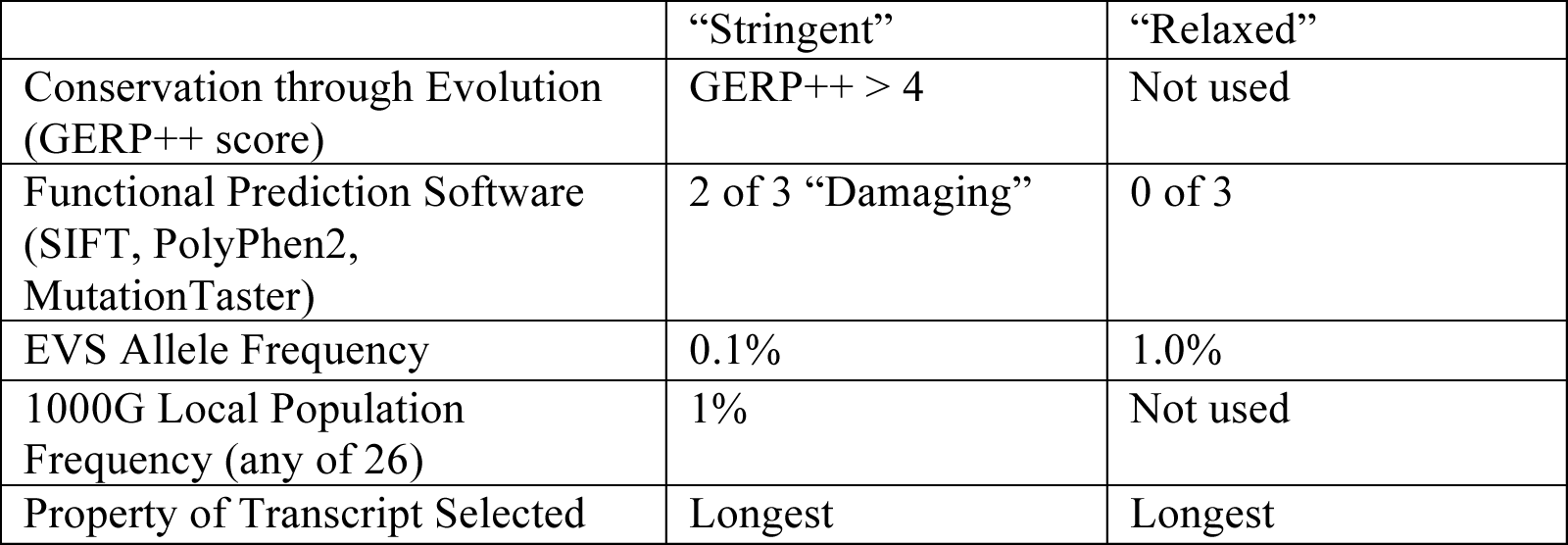
Bioinformatics parameters used for “stringent” or “relaxed” threshold filtering

|  | “Stringent” | “Relaxed” |
| --- | --- | --- |
| Conservation through Evolution (GERP++ score) | GERP++ > 4 | Not used |
| Functional Prediction Software (SIFT, PolyPhen2, MutationTaster) | 2 of 3 “Damaging” | 0 of 3 |
| EVS Allele Frequency | 0.1% | 1.0% |
| 1000G Local Population Frequency (any of 26) | 1% | Not used |
| Property of Transcript Selected | Longest | Longest |

Qualifying variants among these 21 genes were determined as follows: A loss-of-function variant had to have an allele frequency < 0.1% or < 1% in the stringent and relaxed pipelines, respectively. For missense variants, the stringent pathogenicity pipeline consisted of the following: (1) allele frequency < 0.1% in the EVS dataset in each population and <1% in any of the 26 1000G local populations. (2) predicted as damaging by 2 of 3 of the functional prediction algorithms SIFT [29], PolyPhen2 [30], and MutationTaster [31] as annotated in dbNSFP [32]; (3) minimum genomic evolutionary rate profiling++ (GERP++) [33] score of 4; and (4) present in the longest transcript. The “relaxed” pipeline for missense variants consisted of the following: (1) allele frequency < 1% in EVS server, with allele frequency based on 1000G local population not applied; (2) no requirement for predictions as damaging with functional prediction algorithms; (3) GERP++ was not applied; and (4) present in the longest transcript. Variants with allele frequency > 5% in either of the cohorts, but less than 1% in EVS, were attributed to technical artifact from the microfluidic PCR or NGS steps of the pipeline and excluded in all the analyses. Finally, we forced the inclusion of *NPHS2*p.229Q as a qualifying variant whenever it was present because of its known role in monogenic NS, despite it being present at an AF >1% in EVS.

### Variant Analyses

We performed the following analyses comparing frequencies between NS cases (NEPTUNE) and the reference population (1000G), under both the “relaxed” and “stringent” filters.

1. **Prevalence of individuals with qualifying variants per cohort**: To determine overall qualifying variant burden, we calculated the number of individuals with at least one qualifying variant present in any of the genes sequenced.
2. **Prevalence of qualifying variants per person per cohort:** To determine the per-person variant burden, we calculated the number of qualifying variants per person across all genes.
3. **Prevalence of qualifying variants per gene per cohort:** To determine per-gene variant burden, we calculated the total qualifying variants per gene.
4. **Pattern of variants across genes per person:** For those individuals harboring >1 qualifying variant, we examined whether there were combinations of genes harboring variants that were enriched in cases vs reference population.

Sanger confirmation was performed on variants found under the stringent filter only. Parental DNA was not available as part of either study, which precluded ability to perform segregation studies.

### Statistical Analyses

Categorical variables were compared using two-tailed Fisher’s exact test with confidence p=0.05.

## RESULTS

### Per-cohort burden

We first computed the prevalence of individuals with at least one qualifying variant in NEPTUNE and 1000G (Figure 1). Under the stringent filter 16.2% of NS cases and 15.4% of 1000G had at least one qualifying variant (p= 0.74). Using the relaxed filter, 64.7% of cases and 68% of 1000G had at least one qualifying variant (p= 0.24). In total, NS cases did not have an increased burden of heterozygous protein-altering variants in this monogenic NS gene-set. Identified variants including genomic coordinate (using UCSC genome build hg19), reference and alternative allele, variant effect (and amino acid change if non-synonymous coding variant) are listed in **Supplemental Tables 1-4**.

**Figure 1.**
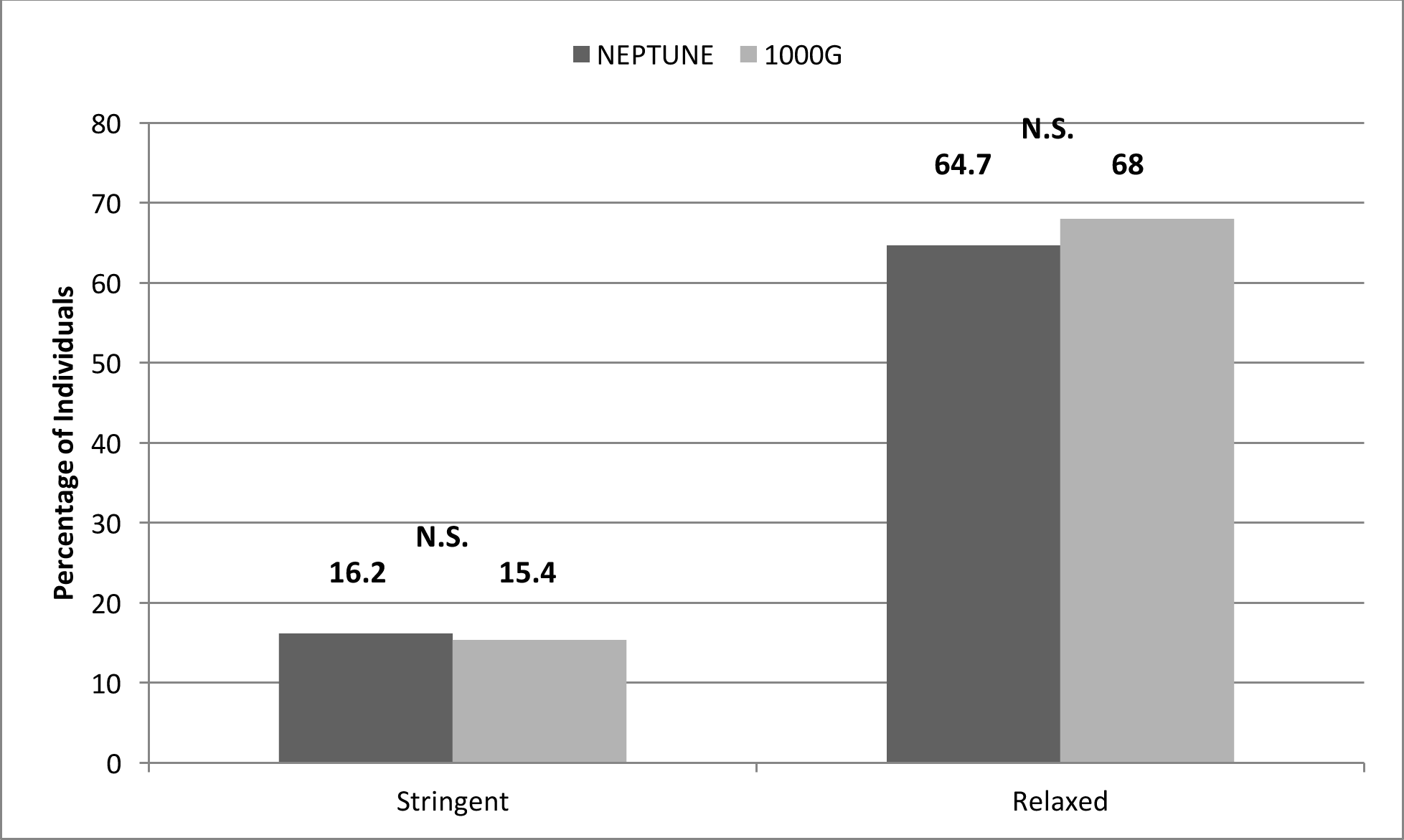
Prevalence of individuals with at least one qualifying variant in NS cases (NEPTUNE) and a reference population (1000 Genomes Project [1000G]). There was no significant difference between cohorts using stringent filter (p= 0.74) or relaxed filter (p=0.24). N.S. denotes no significant difference.

### Per-person burden

We next characterized the distribution of variant burden per person in the NEPTUNE and 1000G (“per-individual burden”) (Figure 2). Using the stringent filter, there was no significant difference between cases and the reference population when comparing individuals with ≥1 qualifying variants (p=0.74), or ≥ 2 qualifying variants (p=0.77). As expected, the percentage of individuals with ≥ 1 qualifying variant increased under the relaxed filter. Under the relaxed filter, there was no significant difference in per-individual burden between cohorts for those with ≥ 2 variants (p=0.07), ≥ 3 variants (p=0.16), ≥ 4 variants (p=1), and > 5 variants (p=0.79) (**Supplemental Figure 1**).

**Figure 2.**
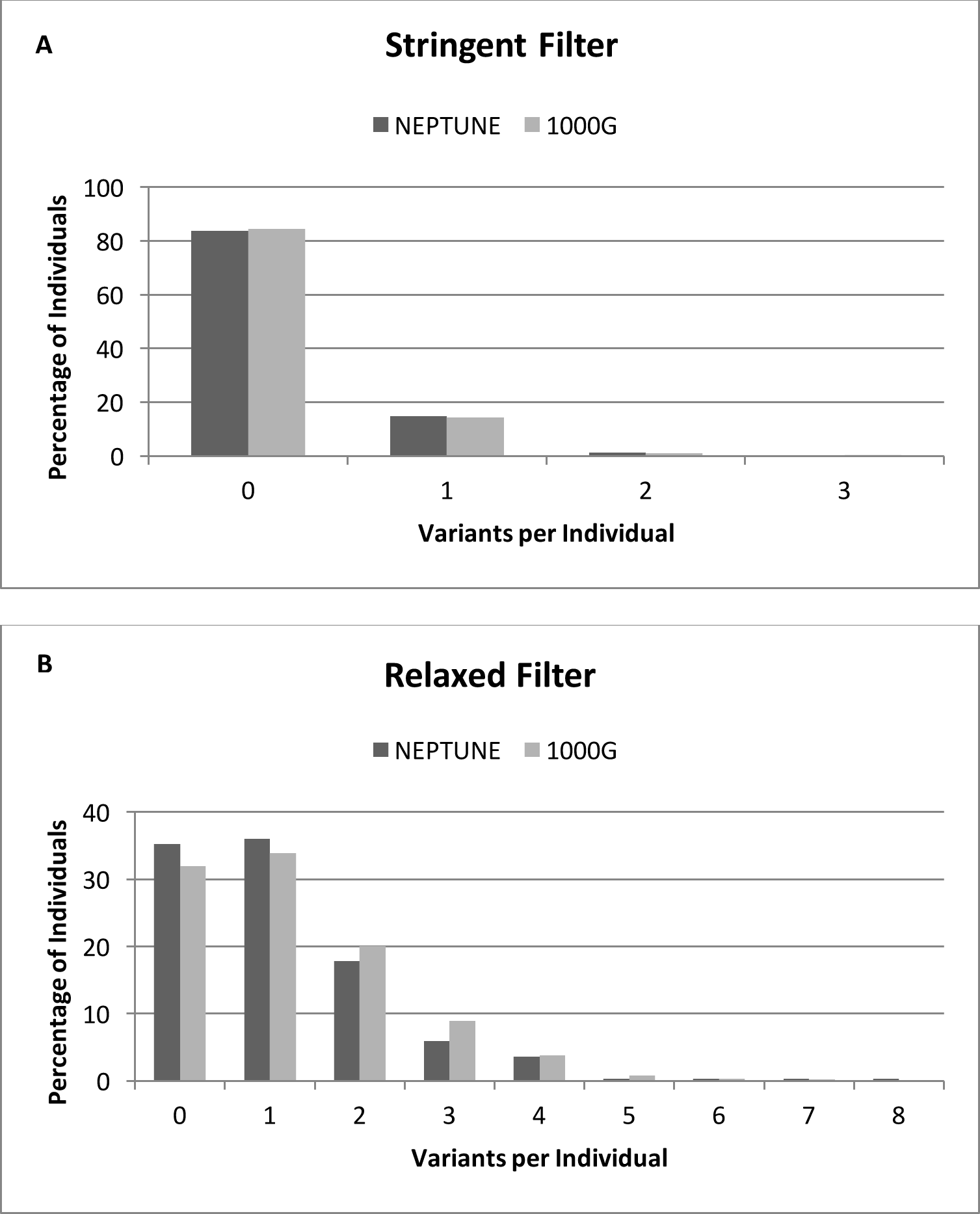
Percentage of individuals with qualifying variants, as compared between NS cases (NEPTUNE) and reference population (1000G). (**A**) Stringent filter. (**B**) Relaxed filter.

### Per-gene burden

We computed the number of qualifying variant per gene in NS cases and the reference population under stringent and relaxed filters (Figure 3). Variant burden per gene was calculated as percentage of total variant burden across all genes. It is known that independent of an individual’s disease status, longer genes have a greater number of variants [34]. This helps explain the larger number of variants seen in longer genes such as *CUBN, PLCE1,*and *PTPRO*in both NEPTUNE and 1000G. Under the stringent threshold, variants were distributed across 12 genes in NS cases and 18 genes in the reference population. The only significant difference in the distribution between cases and the reference population was with *NPHS2*, primarily given inclusion of p.R229Q variant, which is further discussed below. Under the relaxed threshold, variants were distributed between 20 genes in NS cases and 21 genes in the reference population. The burden of variants in *COQ2* (0.3% vs 3.4%; p<0.001), and *PDSS2*(5% vs 8.5%, p=0.02) were significantly higher in reference population, whereas burden of variants in *CUBN* (30.2% vs 21.4%; p<0.001) and *SCARB2* (2.9% vs 1.4%, p=0.04) was higher in NS cases. Of the 103 total variants in *CUBN*identified in NS patients, 16 variants (representing 4 unique alleles: rs140806389, rs78201384, rs117400821, rs191787640) demonstrated an allele frequency (AF) > 1% in the East Asian (EAS) sub-population of Exome Aggregation Consortium (ExAC). An additional 3 variants (rs77886913, rs150985118, rs529907907) demonstrated AF of <1% but the AF in EAS sub-population was higher than in any other sub-population.

**Figure 3.**
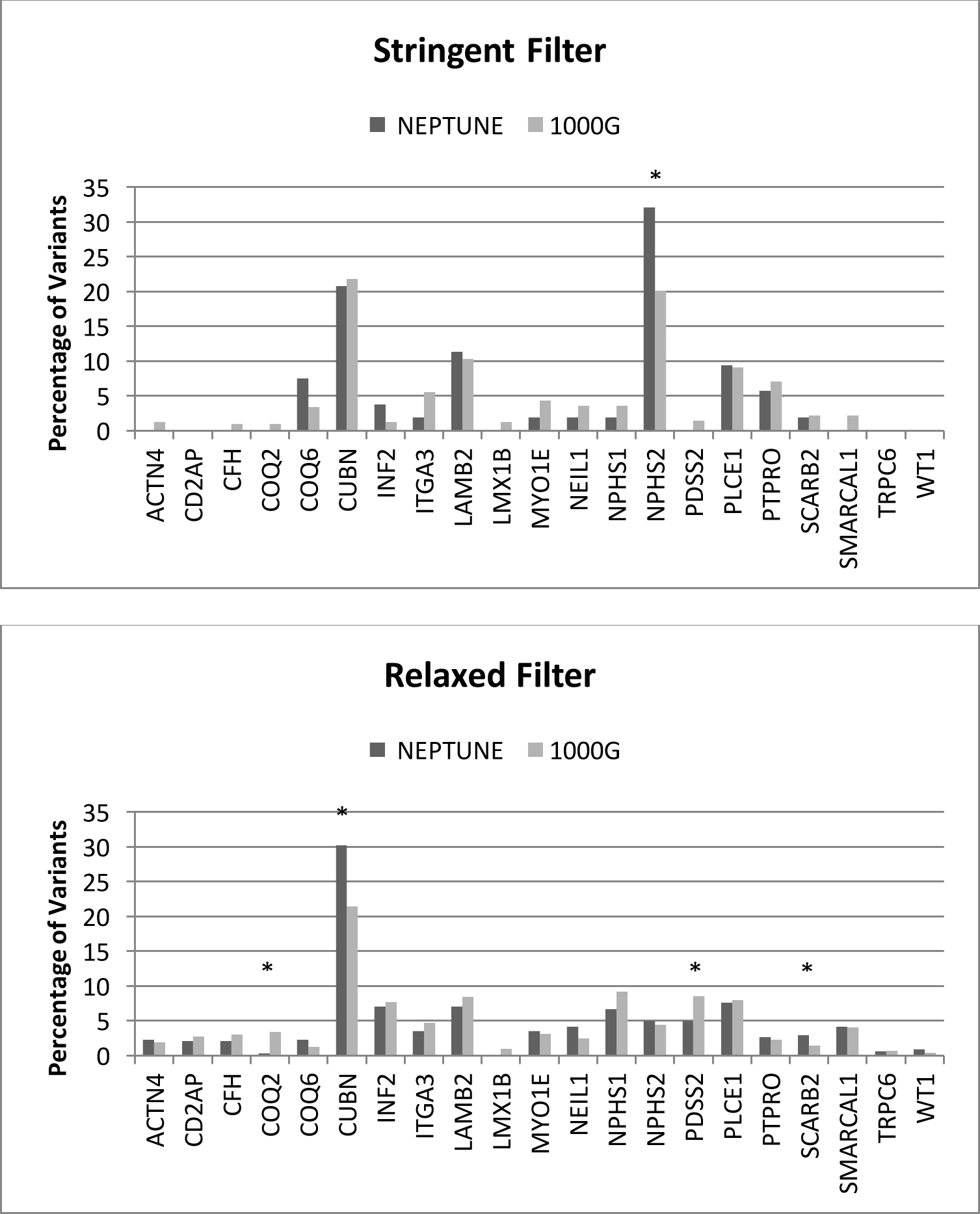
Variants per gene, expressed as percentage of total qualifying variants, between NS cases (NEPTUNE) and reference population (1000G). (**A**) Stringent filter. There was significant increase in *NPHS2* variants in cases, primarily p.R229Q. (**B**) Relaxed filter. Under relaxed filter, variants in *COQ2* and *PDSS2* were significantly higher in 1000G cohort (p < 0.05) while qualifying *CUBN* and *SCARB2* variants were higher in cases. Significant differences are denoted by * symbol.

### Oligogenicity

NEPTUNE patients did not have an enrichment of variants across >1 gene as compared to 1000G subjects. Furthermore, there were no specific combinations of variants across >1 gene that were enriched in NS cases versus the reference population. This was true under the stringent and relaxed filter.

### Inclusion of *NPHS2*, p.R229Q

Given its previous implication in Mendelian disease despite its common frequency in the European population (3%), we forced inclusion of *NPHS2*p.R229Q allele in each of our analyses. In NEPTUNE, 16 individuals heterozygous for the *NPHS2* p.R229Q variant were identified (allele frequency=2.6%). In 14 of these individuals, the p.R229Q variant was the only identified qualifying variant. In the remaining two individuals, one had a single variant in *PLCE1* and the other had a single variant in *LAMB2*. In 1000G, 66 individuals were heterozygous for *NPHS2* p.R229Q variant and 3 individuals were homozygous (allele frequency=1.4%). In 63 of heterozygous individuals, the p.R229Q variant was the only variant identified. No individuals had the p.R229Q variant and another *NPHS2* variant. In the five remaining individuals, two had a single variant identified in *CUBN,*one individual with a single variant in *LAMB2,*one individual with a single variant in *PDSS2,*and one individual with a single variant in *ITGA3*. The higher percentage in NS cases reflects a higher percentage of European individuals in this cohort as compared to 1000G, as 144 of the 303 (47.5%) of NEPTUNE cases were of European ancestry, as compared to 502 of 2498 (20%) of individuals in 1000G. More importantly, in both groups, individuals were likely to only harbor p.R229Q without additional variants in *NPHS2* or other genes, decreasing the likelihood of *NPHS2* p.R229Q contributing to oligogenicity.

### Subgroup analysis of variant burden

Variant burden analysis was performed using subgroups categorized according to pediatric disease onset, history of complete remission, and/ or histologic diagnosis **(Table 5).** A higher percentage of individuals in NEPTUNE cohort who achieved complete remission had at least one qualifying variants, as compared to those who did not achieve complete remission (p<0.01). Using relaxed filter, more pediatric MCD patients than FSGS patients had at least one qualifying variant (p<0.01). When examining either pediatric patients alone or all patients together (adult and pediatric) with either minimal change disease or FSGS, there was no difference using either the stringent or relaxed filter. Finally, under relaxed filtering, 1000G had a higher prevalence of individuals with at least one variant than cohort of pediatric NS patients (p=0.04) or pediatric FSGS patients (p<0.01), but not pediatric MCD patients (p=0.14).

**Table 5.** Analysis of burden of at least one variant in any of the 21 monogenic NS genes in clinically defined subgroup of participants. Statistical analysis was performed using two-tailed Fisher’s exact test. Abbreviations are as follows: Ped = Pediatric, MCD= minimal change disease, FSGS= focal segmental glomerulosclerosis, CR = Complete Remission.

|  |  |  | p-value |
| --- | --- | --- | --- |
| **Pediatric NS vs 1000G** | **Ped NS (n=98)** | **1000G (n=2498)** |  |
| Stringent Filter | 12 (12%) | 384 (15%) | 0.48 |
| Relaxed Filter | 57 (58%) | 1699 (68%) | 0.04 |
| **Pediatric FSGS vs 1000G** | **Ped FSGS (n=30)** | **1000G (n=2498)** |  |
| Stringent Filter | 2 (7%) | 384 (15%) | 0.3 |
| Relaxed Filter | 14 (47%) | 1699 (68%) | < 0.01 |
| **Pediatric MCD vs 1000G** | **Ped MCD (n=44)** | **1000G (n=2498)** |  |
| Stringent Filter | 8 (18%) | 384 (15%) | 0.53 |
| Relaxed Filter | 35 (80%) | 1699 (68%) | 0.14 |
| **Pediatric Complete Remission History** | **Yes CR (n=63)** | **No CR (n=35)** |  |
| Stringent Filter | 8 (67%) | 4 (33%) | 0.22 |
| Relaxed Filter | 38 (66%) | 20 (33%) | 0.01 |
| **Pediatric FSGS vs Pediatric MCD** | **Pediatric FSGS (n=30)** | **Pediatric MCD (n=44)** |  |
| Stringent Filter | 2 (7%) | 8 (18%) | 0.19 |
| Relaxed Filter | 14 (47%) | 35 (80%) | < 0.01 |
| **FSGS+MCD vs 1000G** | **FSGS+MCD (n=162)** | **1000G (n=2498)** |  |
| Stringent Filter | 23 (14%) | 384 (15%) | 0.82 |
| Relaxed Filter | 108 (58%) | 1699 (68%) | 0.73 |
| **FSGS + MCD (Ped) vs 1000G** | **FSGS+MCD (n=74)** | **1000G (n=2498)** |  |
| Stringent Filter | 10 (14%) | 384 (15%) | 0.75 |
| Relaxed Filter | 49 (58%) | 1699 (68%) | 0.8 |

## DISCUSSION

Motivated by its biologic premise, reports from animal models [35], human sequencing case reports [15, 16, 18, 19], and our previous finding that found only 2.9% of US patients with NS had a monogenic form of disease [14], we hypothesized that a proportion of the heritability of NS may result from protein-altering risk alleles within Mendelian NS genes, acting alone or with an oligogenic architecture. To test this hypothesis, we compared the prevalence of putative risk alleles and oligogenicity in an unselected NS case cohort recruited from the United States and a worldwide reference population. When using this strategy under a stringent pathogenicity filter, we found NS patients did not have an increased burden of risk alleles in these genes; nor was there any evidence of oligogenicity. Specifically, the NS cohort did not have a greater overall prevalence of qualifying variants. NS patients neither had a greater average number of qualifying variants, nor were there a group of outliers with increased variant burden. From a gene perspective under a stringent filter, no specific gene had a greater burden of qualifying variants in cases vs the reference population, while population stratification is implicated in the enrichment of *CUBN* and *SCARB2* variants under a relaxed filter. Lastly, there were no specific variant-harboring gene pairs enriched in NS cases.

We also created a “relaxed” filter in order to determine whether variants in this geneset, predicted to be less pathogenic, could nonetheless explain some of the heritability of NS. While more frequent in the population or less evolutionarily conserved, we hypothesized that these low-frequency protein-altering variants could nonetheless act as hypomorphic alleles and deleteriously impact the function of one or more proteins. As mentioned before, this form of genetic architecture has been illustrated with the p.R229Q variant of podocin in NS patients of European ancestry [23, 36]. Yet, under the relaxed filter as well, we also did not identify any significant differences in variant burden between NS cases and the reference population. For the gene-level burden analysis, 2/21 genes demonstrated higher variant burden in cases whereas 2/21 genes demonstrated higher variant burden in the reference population. We believe that this enrichment is reflective of population stratification, with alleles that are rare in the African and European subjects in the EVS nonetheless being more common in Asian populations of the 1000G cohort. This population stratification under the relaxed filter was specifically interrogated for qualifying variants in *CUBN*. These specific findings support that population stratification is a potential confounder in the interpretation of variants and must be properly accounted for in sequence analysis of any NS patient in research or clinical care.

The negative results from our study here differ from previous reports of oligogenicity in NS. There are a number of potential explanations for this discordance. One possibility is that in previously reported cases, the presence of heterozygous or “third” alleles purported to contribute to the pathogenesis of NS were present coincidentally and in fact non-contributory to causing NS or modifying its phenotype. Here we found that 15.4% of a reference population harbors a variant in a monogenic NS gene (a putative risk allele) under stringent criteria and this increased to 68% when using a relaxed criteria (in which variants still needed to have an allele frequency <1%). Furthermore, 33% of the reference population had 2 or more qualifying variants under relaxed thresholds. Given the lack of standardization of pathogenicity prediction algorithms, particularly in the research setting, there is substantial risk of misclassifying benign or background protein altering variants in NS genes as deleterious depending on the stringency of the algorithm.

Another possibility is that our bioinformatics-based filtering strategy is insufficiently accurate to differentiate pathogenic variants from those that are benign. We do not have parental data which would help with segregation of alleles in cases and the reference population. In addition, with more accurate filters, harmless variants would drop out and we could hypothesize that cases may be enriched for pathogenic variants. Perhaps more sensitively than bioinformatics filters, functional testing of variants discovered in sequencing studies could help identify specific variants that damage the function of the protein product. Studies in Type 2 Diabetes Mellitus, for instance, have shown the benefit of this strategy in identifying *bona fide* disease associated rare variants [37].

A final possibility is that our current study is underpowered to detect small but significant enrichment of risk alleles and oligogenicity within known NS genes in NS cases as compared to those without disease. Future studies in large independent cohorts and/or meta-analyses would be one strategy to empower us to discover these small effects. Ultimately, we suggest that similar studies should be performed in larger, independent cohorts of NS cases and controls to determine the generalizability of our discovery. This should be feasible given the number of cohorts that have already generated targeted sequence data of these genes for monogenic disease prevalence studies.

We suspect that the answer likely lies somewhere in the middle. As already illustrated by the low frequency p.R229Q variant of *NPHS2* and its dependency of specific mutations on the *trans* chromosome [36], there may be other complex forms of genetic variation in known Mendelian NS genes that cause NS, or greatly increase its risk. Future studies of thousands of NS cases may ultimately identify additional protein-altering risk alleles for NS in these specific genes or others, exome-wide. But in many other cases, particularly when using insufficiently stringent filtering strategies, standing background variation in monogenic NS genes is likely misclassified as pathogenic, and is neither contributing to the cause of NS nor modifying its phenotype.

As we continue to sequence increasing numbers of implicated NS genes in increasing numbers of affected patients, we are pressed to maximize our accuracy in classifying our patients with genetic forms of NS. This is necessary for the genetic information to be clinically meaningful in guiding prognosis, treatment and counseling decisions. Here, we determined that while NS patients harbored combinations of protein-altering alleles across known monogenic NS genes, they did so at rates not significantly different than members of a reference population. Moving forward, the development of more advanced bioinformatics filtering, and/or increased use of functional testing of variants should help us identify whether more complex forms of genetic variation in known Mendelian NS genes contribute to, or cause, NS. For now, however, particularly in the absence of strongly supportive functional information or complementary genetic support, we suggest using caution in attributing a pathogenic role to non-causal variants identified in implicated NS genes.

## ACKNOWLEDGEMENTS

Brendan Crawford is supported by the National Institutes of Health Kidney Research Training Grant T32DK007378.

Matthew Sampson is supported by the Charles Woodson Clinical Research Fund, RO1DK108805, and is a Carl W. Gottschalk Research Scholar of ASN Foundation for Kidney Research.

The Nephrotic Syndrome Study Network Consortium (NEPTUNE), U54-DK-083912, is a part of the National Institutes of Health (NIH) Rare Disease Clinical Research Network (RDCRN), supported through a collaboration between the Office of Rare Diseases Research (ORDR), NCATS, and the National Institute of Diabetes, Digestive, and Kidney Diseases. Additional funding and/or programmatic support for this project has also been provided by the University of Michigan, the NephCure Kidney International and the Halpin Foundation.

